# Swarming bacteria exhibit developmental phase transitions to establish scattered colonies in new regions

**DOI:** 10.1101/2024.09.24.614802

**Authors:** Amanda M. Zdimal, Giacomo Di Dio, Wanxiang Liu, Tanya Aftab, Taryn Collins, Remy Colin, Abhishek Shrivastava

## Abstract

The bacterial Type 9 Secretion System (T9SS) is essential for the development of periodontal diseases and Bacteroidetes gliding motility. T9SS-driven motile bacteria, abundant within the human oral microbiota, transport non-motile oral microbes and bacteriophages as cargo, shaping the spatial structure of polymicrobial communities. However, the physical rules governing the dispersal of T9SS-driven bacterial swarms are barely understood. Here, we collected time-lapse images, under anaerobic conditions, of developing swarms of a T9SS-driven microbe common to the human oral microbiota. Tracking of swarms revealed that small peripheral flares emerging from a colony develop structures that resemble fireworks displaying a chrysanthemum effect and flower-like patterns that convert to wave-like patterns and which further evolve into scattered microcolonies. Particle-image velocimetry showed density-dependent phase transitions and initial vorticity within these emerging patterns. Numerical simulations demonstrate that these patterns arise due to changes in swarm speed and alignment strength. Our data reveal a strategy used by an anaerobic swarming bacterium to control swarm behavior, resulting in scattered microcolonies distant from the mother colony, thus reducing competition for resources among colony members. This might ensure species survival even if conditions change drastically in one location of the human oral cavity.

## INTRODUCTION

Collective motion is common in biology, on both microscopic and macroscopic levels (1, 2). The widespread use of coordinated movements amongst members of a group implies evolutionary benefits of these behaviors (3). Motility is imperative in the search for favorable environmental conditions and scavenging for water and adequate food sources. The ability of many members of a group to mobilize ensures survival of the species from predators, waste accumulation and environmental challenges. Although motility and collective motion are found to often provide clear benefits, the mechanisms controlling these essential behaviors in bacteria are poorly understood.

Motile bacteria serve as excellent models for studying the mechanics of motility and collective motion in their simplest forms. Bacteria have developed various strategies for movement, highlighting the significance of mobility across all scales of life. Most bacteria rely on extracellular appendages to facilitate movement; for instance, type IV pili enable twitching motility (4), and flagella allow bacteria to swim through liquids and swarm across surfaces. The well-characterized flagellar motility is driven by proton motive force (pmf) and it is arguably the most common type of motility utilized by bacteria (5, 6). Another recently discovered motor that drives motility is the Type 9 secretion system (T9SS), which transports essential proteins across the outer membrane to execute gliding motility (7). T9SS is one of the three known pmf-driven biological rotary motors, the other two being ATP synthase and the bacterial flagellar motor (8, 9). In the T9SS system, a rotary motor powered by proton motive force drives the transport of cell surface adhesins across the surface in helical fashion (10, 11). The adhesins attach to a helical track on the cell surface and interact with a boundary such that the movement of the adhesin across the cell surface propels the cell forward (12, 13). Gliding machinery is common across the Bacteroidetes phylum, but not all T9SS-containing organisms are capable of motility.

Bacteria of the genus *Capnocytophaga*, typically isolated from the human oral microbiota, are driven by T9SS and serve as potential models for studying gliding motility and its impact on host-associated polymicrobial communities. *Capnocytophaga* sp. coordinate gliding motility actions between cells to result in collective motion through a process known as swarming (14–16). Interestingly, *Capnocytophaga* is the only genus in the human oral cavity that utilizes T9SS-driven gliding motility, marking it as an important representative of the Bacteroidetes phylum in the human oral microbiota (17). The gliding motility and T9SS of *Capnocytophaga ochracea* is crucial for the formation of single-species biofilms (18). The genus *Capnoctophaga* is one of the most prominent bacterial genera in the oral microbiome, reaching colonization levels up to 8% of the total bacteria present in the supra-gingival and sub-gingival plaques of healthy humans (19). Studies aimed at characterizing the role of *Capnocytophaga* sp. in the oral cavity are minimal but have shown its ability to swarm and carry other bacteria (20), as well as bacteriophages (21) as hitchhikers. *Capnocytophaga* sp. are detected in plaque deposits tenfold higher than in non-plaque sites in healthy individuals (19). Additionally, it is also considered as an opportunistic human pathogen (22, 23). Taken together, *Capnocytophaga sp.* appears to play a complex and crucial role in shaping the structure of the human oral microbiota and overall human health.

One challenge in studying swarm development of *Capnocytophaga* sp. is that it requires anaerobic conditions for growth and motility. Bacteria currently used as models for swarming behavior thrive in aerobic conditions, which allows for easy observation of swarm development and structure (24–29). Many of these studies have shown that bacteria display unique phenotypes with different swarm sizes and patterns. Swarms of aerobic bacteria have been analyzed for properties that include pattern formation, speed, vorticity, and the direction of movement. Due to the interest in characterizing swarm development, many models have been developed that aim to facilitate our understanding of swarm processes (30–38). These models demonstrate the phase separation of particles (39) and multiple studies have confirmed the presence of phase transitions in bacterial swarms that align with numerical simulations (40–42). However, to our knowledge, no study has used an anaerobic swarming bacterium as a model for collective motion.

We investigated motility and swarm development of *C. ochracea* under anaerobic conditions. Surface stiffness and nutrient composition were altered to assess how these factors influence swarm capabilities of *C. ochracea.* We quantified changes in swarm size and patterns as a function of varying agar concentration and nutrient composition. Long-term imaging revealed that *C. ochracea* swarms exhibit striking spatiotemporal patterns reminiscent of fireworks displaying a chrysanthemum effect, characterized by circular forms with jagged edges. Later, one half of the flower-like structure expands outwards, transitioning into wave-like patterns. Timelapses of developing *C. ochracea* swarms were captured, and their movements were tracked using a modified version of classical particle image velocimetry (PIV). This analysis uncovered phase transitions dependent on cell density and alignment, providing physical insights into swarm patterning and behavior. Toward the end, the swarms coalesced to form scattered microcolonies, which were millimeters to centimeters away from the mother colony. This behavior reveals a mechanism that could be akin to seed dispersal in plants or spore dispersal in fungi, potentially increasing the survival probability of swarming bacterial species under adverse conditions.

## MATERIALS AND METHODS

### Bacterial strains, media and growth conditions

*C. ochracea* (ATCC 27872) was cultured on Trypticase Soy (Becton Dickinson) agar plates supplemented with 3 g/L yeast extract and 5 μg/mL Hemin (TSY). The plates used in this study had varying concentrations of Bacto agar (Becton Dickinson), between 0.5% and 2%. To investigate how blood changes swarming, 5% defibrinated horse blood was added to a subset of the plates (TSY-blood). *C. ochracea* was maintained on TSY with 1.5% agar, in an anaerobic chamber at 37°C with a final gas mixture of 80% N_2_/17.5% CO_2_/2.5% H_2_. For swarm plate assays, petri plates were incubated in anaerobic boxes (Becton Dickinson) with a high CO_2_-producing anaerobic sachet (Mitsubishi) and four small flasks containing ∼30 mL H_2_O each to maintain a high relative humidity. *C. ochracea* τι*gldK* strain, a gift from Kazuyuki Ishihara from Tokyo Dental College, was used as a control for motility (18).

### Swarm assays on varying surface stiffness and nutrient composition

Swarm assays were performed on TSY plates with and without blood, and with 0.5%, 1%, 1.5% or 2% agar. Plates were made fresh on day 0 and allowed to set for 30 min before inoculation. A 2-day old *C. ochracea* swarm plate was scraped in 1 mL TSY broth, and the OD_600_ was adjusted to 0.8. A volume of 5 μL of the culture was spotted on fresh plates and air dried in a laminar flow biosafety cabinet for 30 min. Plates were then transferred to an anaerobe box containing water flasks and incubated under high CO_2_ for 7 days. Swarm size was measured using swarms inoculated on TSY plates, and images were acquired using a stereoscope (AmScope, MU1809) on days 1, 2, 3, 5 and 7. Four biological replicates were acquired. To improve the contrast of images for the **Figure 1**, 0.05% black food coloring (Chefmaster liqua-gel coal black food color, Fullerton, CA) was added to the agar (dyed TSY). A preliminary experiment was performed using regular TSY and dyed TSY plates to ensure the dye had no effect on *C. ochracea* swarming **(Figure S1)**.

**Figure 1.**
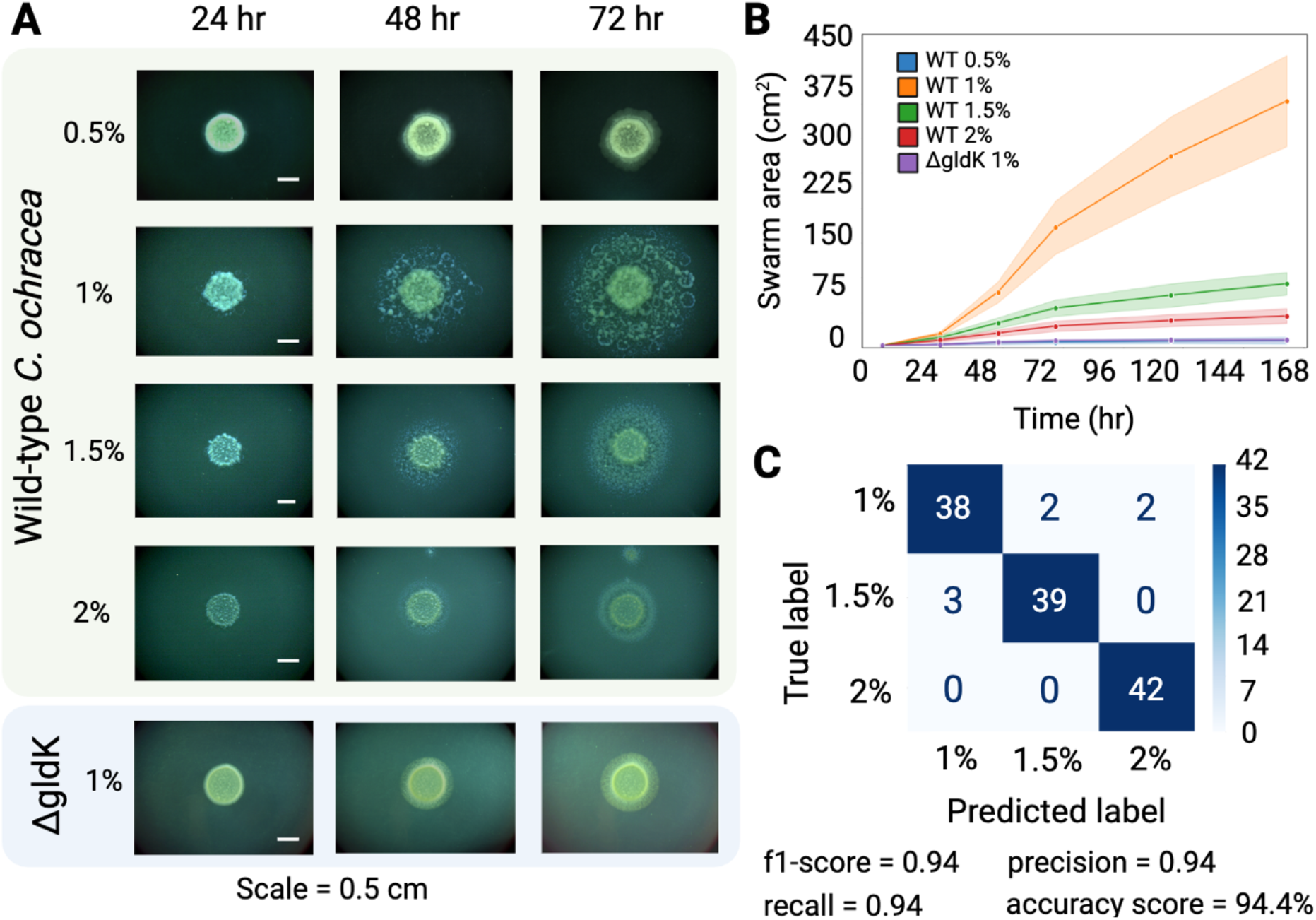
*C. ochracea* swarms exhibit distinct phenotypes on surfaces of varying stiffness. Cell suspensions spotted on TSY agar plates were incubated anaerobically for 7 days. Images were taken at 24, 48, 72, 96 and 168 h. **(A)** Images of wild-type swarms demonstrate that different swarm patterns and sizes result from varying surface stiffness. The T9SS mutant *τιgldK* is used as a motility-deficient control. **(B)** Swarm areas throughout the course of development indicate that swarms on 1% agar are significantly larger than swarms on the other agar concentrations. **(C)** An AI-based algorithm trained on a subset of swarm images from TSY plates with 1%, 1.5% and 2% agar differentiated the test images with around 94% accuracy, thus showing that the swarm patterns are reproducible. Colorbar shows the number of images tested by the trained algorithm.

### AI-based evaluation of swarm patterns on different agar concentrations

A Support Vector Machine (SVM) classifier algorithm was used to determine whether the distinct swarm patterns on surfaces of varying stiffness were a reproducible phenomenon. Sixty swarm plates on dyed TSY were prepared for each agar concentration: 1%, 1.5%, and 2%. A total of 180 images, taken at 72 hours, were used in the study. Out of the 60 images per agar concentration, 18 (30%) were used for training, and 42 (70%) were analyzed by the algorithm. The algorithm then predicted the agar concentration used in the images based on the training phenotypes.

### Swarm timelapse acquisition

*C. ochracea* swarm development was investigated using a stereoscope over the course of 5 days. TSY with 1% and 1.5% agar were used for swarm tracking. Plates were inoculated as described above, then transferred to a humidified anaerobe box fixed to a stereoscope using double-sided tape, and housed in a 37°C incubator. Timelapses were started at 20 hours to allow sufficient time for the condensation produced by the anaerobe sachets to dissipate so that clear images could be obtained. Images were acquired every 5 min for ∼3-4 days, until the swarm front exceeded the field of view. To ease the burden of image capture and analysis, pixels were binned (4X) to produce images of 1228 x 920 pixels. Timelapses were repeated in triplicate.

### Particle image velocimetry tracking of developing swarms

Movies were first preprocessed in ImageJ to even out the non-constant illumination profile and to reduce noise. For this, a background image obtained from averaging 50 images of virgin agar plates was subtracted from all images in the movie. Next, the [I_min_ ; I_max_] integer grayscale of the background subtracted images was recast on a 8-bit gray scale ([0; 255]). Finally, a filter was applied to improve signal over noise ratio (S/N). A 3D median filter (σ_X_ = 0 px, σ_Y_ = 0 px, σ_t_ = 3 frames) was applied to the 1% agar movies, but we had to apply a stronger 3D mean filter (σ_X_ = 2 px, σ_Y_ = 2 px, σ_t_ = 3 frames) to the 1.5% agar movies, because the stronger light scattering at this concentration resulted in lower S/N.

A previously developed Particle Image Velocimetry algorithm (43) was then applied to the movie to measure local velocities. In short, a square window of size *a* = 64 px is extracted around each interrogation point on a rectangular grid (spacing *da* = 8 px for 1% agar movies, 4 px for 1.5% agar) and blurred with a Gaussian profile (exp(-*r*^2^/*l*^2^), *l* = 7 px for 1% agar movies, 10 px for 1.5% agar) centered on the interrogation point, to improve spatial resolution. The local cumulative drift ***r_k_***(*t*) was measured using a Fourier transform based algorithm on the blurred interrogation window (43, 44), by fitting the phase shift Δφ**_k_** of its spatial Fourier components as Δφ**_k_**(***q***,*t*) = ***q.r_k_***(*t*)+noise, within the range of wave vectors ***q***=(*q_x_*,*q_y_*) defined by *|q_x_|,|q_y_|< q_max_* = 1.371 px^-1^. The drifts were then temporally median filtered on a 5 frames sliding window, to reduce noise further, and the filtered drifts were linearly fitted, ***r_k_****(t’)=**v***(*t*)(*t’*-*t*)+***r_k_***(*t*), on a 10 frames sliding window to extract the time-resolved local drift velocity ***v****(t)*.

Parameter choices were adjusted to the experimental conditions at hand and obey the following constraints (43, 44): (i) The choices of *a* and *q_max_* reflect the necessity to avoid phase wrapping and to have a good statistics for the fit of the phase, which respectively impose *q_max_|**v**|*<*π* fr^-1^ and *aq_max_*/2*π* ≥ 14. (ii) The choice of *da* trade-offs between computation time and the typical size of the cell motion features, which are smaller on 1.5% agar. The Gaussian blur length *l* is adjusted to capture as much detail of the cell motion as possible while avoiding underestimation of speed. If the blur length *l* is too small, the motion between successive frames of the few-pixels-thick intensity profiles is not fully captured, leading to an underestimation of speed. On the other hand, if the blur length is too large, it results in non-local measurements that account for too large an area around the interrogation point. Therefore, we chose the smallest possible blur length before the typical magnitude starts decreasing.

### Data analysis of PIV timelapses

Custom MATLAB scripts (43, 45) were used to extract velocity maps from the raw PIV data. We then calculated the norm of the velocity, the vorticity and the divergence of microswarmers velocity field using custom Python codes. We computed normalized versions of the vorticity and divergence to dampen the effects of swarmer density being inhomogeneous and focus on gradients of motion directions, 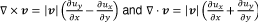, with **u** = **v**/|**v**|. These data were used to characterize temporal and spatial arrangement of *C. ochracea* swarms. Various statistical analyses were performed to quantify swarm development. All data and custom scripts are freely available online on our GITHUB page (https://github.com/amandazdimal/CapnoPIV.git).

### Agent based modeling of swarm behavior

This simulation uses principles of randomization, local interaction (density and alignment), and periodic division to simulate the movement and behavior of cells on a surface. It visualizes how cells interact, align, and move over time, showcasing emergent patterns and behaviors in simulated cellular systems. Individual cells are initially simulated and each cell is initialized with attributes for its position and orientation. Cells are positioned using Cartesian coordinates, offering flexibility in their placement. The initial position (*x_i_*, *y_i_*) of each cell is determined by a randomly generated radius *r_seed_* which is determined by *R* multiplied by the square root of a random number between 0-1. Here, *R* is the radius parameter ensuring uniform distribution within a circle. When generating individual cells around a seed point (eg., due to cell-divison), *R* corresponds to a value of 2. A total of 400 cells are generated per seed and only one seed is used in this simulation. The initial orientation angle of the cell is θ_*i*_, and the speed of cellular motion is *v*.

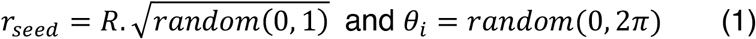

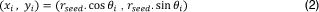

The progression of the simulation is visualized by representing each cell as an oriented rod, which iterates through a series of discrete time steps of 0.1. To reproduce the initial vorticity observed in our experiments an initial ω (rate of change in orientation) of 0.1 is provided in the simulation. The position of the cell changes over time to simulate the motility of the cell as the following:

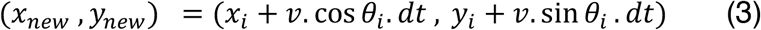

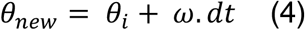

The speed and alignment of each cell are governed by their local environment, with the local density *D*(_*x, y*_) around a cell calculated as follows:

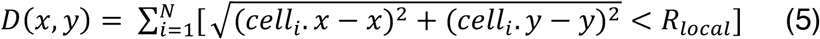

Here *R*_01230_ represents the distance around each cell within which other cells are considered. This defines the scope of the local neighborhood for each cell, and which neighboring cells influence one another. In other words, this determines how far out from a given cell the simulation looks to find neighboring cells. The alignment of a cell is governed by the density threshold, *D_t_* which varies from 10 to 50. and the alignment strength, *A_str_* which varies from 0.07 to 0.21. *D_t_* dictates the cell speed based on local density as follows:

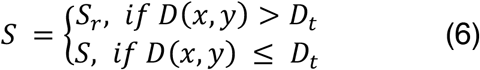

*S* is the speed of the cell, and *S_r_* is the reduced speed of 4 after the cell encounters a density exceeding *D_t_*. *A_str_* determines the degree of alignment with neighboring cells. It influences the orientation of the cell, θ_*new*_. The average angle (θ_*avg*_) is calculated by first averaging the sine and cosine components of neighboring cells, then finally determined by using the Python function arctan2. θ_*i*_ represent the current orientation of the *i^th^* cell, *n* is the number of cells within *R_local_* and Δθ is the shortest angular distance between θ_*avg*_ and θ_*i*_.

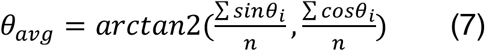

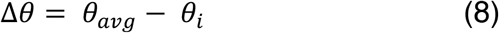

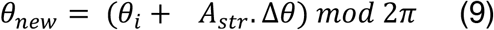

## RESULTS

### *C. ochracea* swarm patterns and sizes change as a function of nutrient composition and surface stiffness

The area and structure of *C. ochracea* swarms were examined on TSY plates with and without blood, and with adjustments to surface stiffness by altering the agar concentration from 0.5% to 2%. On surfaces of optimal stiffness, *C. ochracea* expanded from the edge of the inoculation spot and this expansion ultimately led to a subset of cells becoming immotile and forming microcolonies. The remaining motile cells continued to swirl outward, thus creating distinct motile and nonmotile regions within the swarm.

Swarming was absent on all plates with 0.5% agar, irrespective of the nutrient composition. The colony sizes on these plates were similar to those of the T9SS deficient, non-motile *ΔgldK* strain (**Figure 1A, S2**). Optimal swarming, marked by extensive surface coverage of *C. ochracea* swarms, and a distinctive phenotype characterized by circular and wave-like strings of microcolonies, was observed on plain TSY plates containing 1% agar. Swarms on 1.5% and 2% agar showed patterns similar to one another, although those on 2% agar were smaller and featured tighter, less spread-out microcolonies. On both 1.5% and 2% agar, the microcolonies exhibited more pinpoint or colony-like formations, unlike those observed on plain TSY with 1% agar (**Figure 1A**). Additionally, significant differences in swarm sizes on TSY agar with 1%, 1.5%, and 2% agar were observed (**Figure 1B, Table S1**). TSY blood plates containing 1% agar exhibited a significant reduction in swarm area, nearly 65% smaller compared to plain TSY with 1% agar and displayed considerably fewer microcolonies. Interestingly, unlike the 1% agar conditions, on both 1.5% and 2% agar the sizes and patterns of the swarms remained consistent regardless of the presence or absence of blood (**Figure 1A, B, S3**).

To assess the reproducibility of swarm patterns at different agar concentrations, a Support Vector Machine (SVM) classifier algorithm was employed (46, 47). Images of 72-hour-old *C. ochracea* swarms on 1%, 1.5%, and 2% agar plates were captured, with sixty images for each concentration of agar. Eighteen images (30% of the data) from each concentration were randomly selected to train the classifier, while the remaining forty-two images (70% of the data) served as the test set. As depicted via a confusion matrix, swarms on 2% agar were identified correctly 100% of the time. The identification accuracy for swarms on 1% and 1.5% agar was 90% and 93%, respectively. These findings demonstrate reproducible phenotypes with distinct characteristics on surfaces of identical stiffness for each condition, with an overall accuracy of 94.4%. The precision and recall values, calculated as true positive / (true positive + false positive) and true positive / (true positive + false negative) respectively, were both 0.94 (**Figure 1C**), thus confirming the presence of reproducible patterns in swarm images that change as a function of surface stiffness.

### Tracking spatiotemporal changes in swarm behavior

A live imaging setup (**Figure 2A**) was designed, allowing for the capture of time-lapse images of anaerobic swarms of *C. ochracea* over several days (**Figure 2B**). Approximately 20 hours after inoculation on agar, the structural features of the swarm start to emerge. Therefore, recordings commenced around 20 hours and continued until the swarm front extended beyond the field of view. Images of swarms on varying agar concentrations were captured every 5 min **(Figures 2, S4 and S5, Movies S1-S3, Supplementary text).** The swarm behavior of *C. ochrachea* was quantified using a modified version of traditional Particle Image Velocimetry (PIV), which is based on Phase Differential Microscopy **(Movies S4-S6)**. This approach (see methods) calculates the local velocity field *v*(*r*) by averaging over several neighboring bacteria. Changes in pixel intensity are employed to assess the speed and directionality of swarm development (43). Following PIV, velocity maps were extracted and plotted as heatmaps **(Movies S7-S9)** that demonstrate spatial changes in swarm velocity. Additionally, quiver plots **(Movies S10-S12)**, which display the velocity vectors identified via PIV show the directionality and relative speed of movement of the swarms.

**Figure 2.**
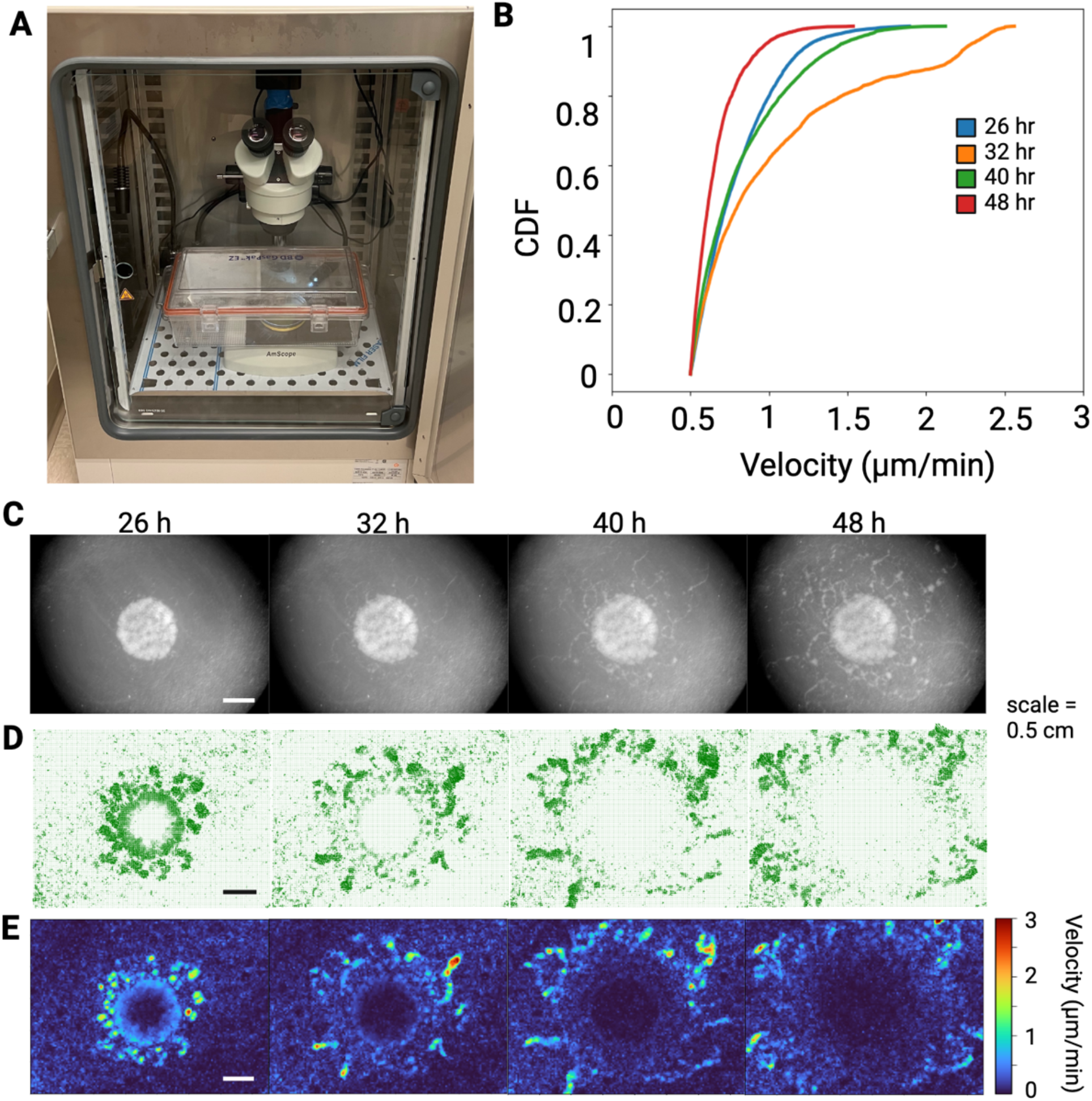
Developing swarms undergo phase transitions. Live imaging of developing swarms were captured over the course of two days. **(A)** Live imaging setup for anaerobic swarms. **(B)** A cumulative density function of swarm speeds during the different phases, showing greatest speeds at 32 h, and slowest speeds at 48 h. **(C)** Stereoscope images of developing swarms from different timepoints. **(D)** Particle image velocimetry of the swarms allowed tracking of swarm behavior and identification of distinct spatiotemporal features as demonstrated by quiver plots. **(E)** Heatmaps of the velocity of developing swarms measured by image velocimetry.

Plotting *v*(*r*) of *C. ochracea* swarms on a two-dimensional image plane provided a spatiotemporal map of swarming. This mapping revealed characteristics that distinguish *C. ochracea* swarms from bacteria that employ other motility mechanisms, such as flagellar-driven swarming and the distinct type of gliding motility of Myxobacteria (48, 49). Approximately 26 hours after inoculation, *C. ochracea* swarms were primarily active around the periphery of the initial inoculation area. Small groups of swarming cells, hereafter referred to as micro-swarmers, emerged from the inoculation region. Interestingly, the micro-swarmers initially appeared as irregularly shaped offshoots disconnected from the parent colony (**Figure 2C, 2E**). However, many micro-swarmers expanded radially, displaying a pattern similar to fireworks bursting with a chrysanthemum effect (**Figure 3A**). This pattern of movement by the microswarmers was characterized by high flow vorticity, i.e. counterclockwise rotational movement, and large divergence, quantifying the expansion from the center of the chrysantemum **(Figure 4A-C)**. These bursting micro-swarmers later transitioned to a flower-like pattern, characterized by circular forms with jagged edges (**Figure 2D, Figure 3C, Figure 3D, Movie S7**). The Cumulative Distribution Function (CDF) displays the arrangement of the *v*(*r*) measured throughout the timelapse. At 26 hours *v*(*r*) shows a range from 0.5 µm/min to 1.5 µm/min, characterized by a long tail, indicating the presence of a subset of faster moving cells within the majority of slower cells.

**Figure 3.**
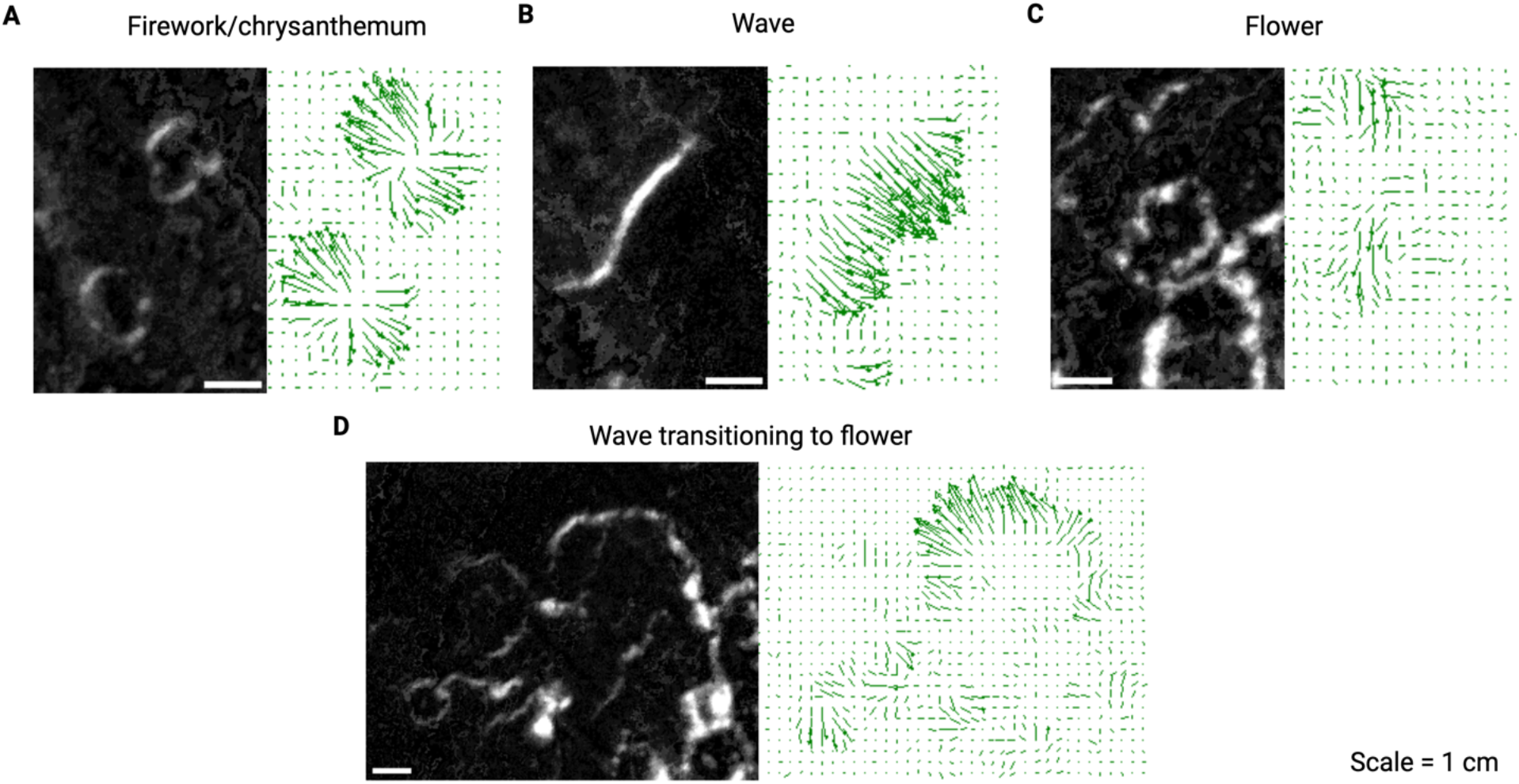
Microswarm expansion patterns. Developing *C. ochracea* swarms exhibit different phases of expansion. The zoomed-in swarm image is accompanied by quiver/velcoity plot of that region. **(A)** Firework or chrysanthemum-like patterns where swarmers expand in all directions from a central point. **(B)** Wave-like movements where swarmers align along a single plane and move in the same direction causing a swarm front. **(C)** Flower-like movements where swarmers move in broken or irregular circular patterns causing a rippled-like expansion similar to flower petals. **(D)** Transition from wave to flower-like expansion, characterized by breaks in the plane of movement, changes to the directionality of some microswarmers, and the emergence of flower-like structures.

**Figure 4.**
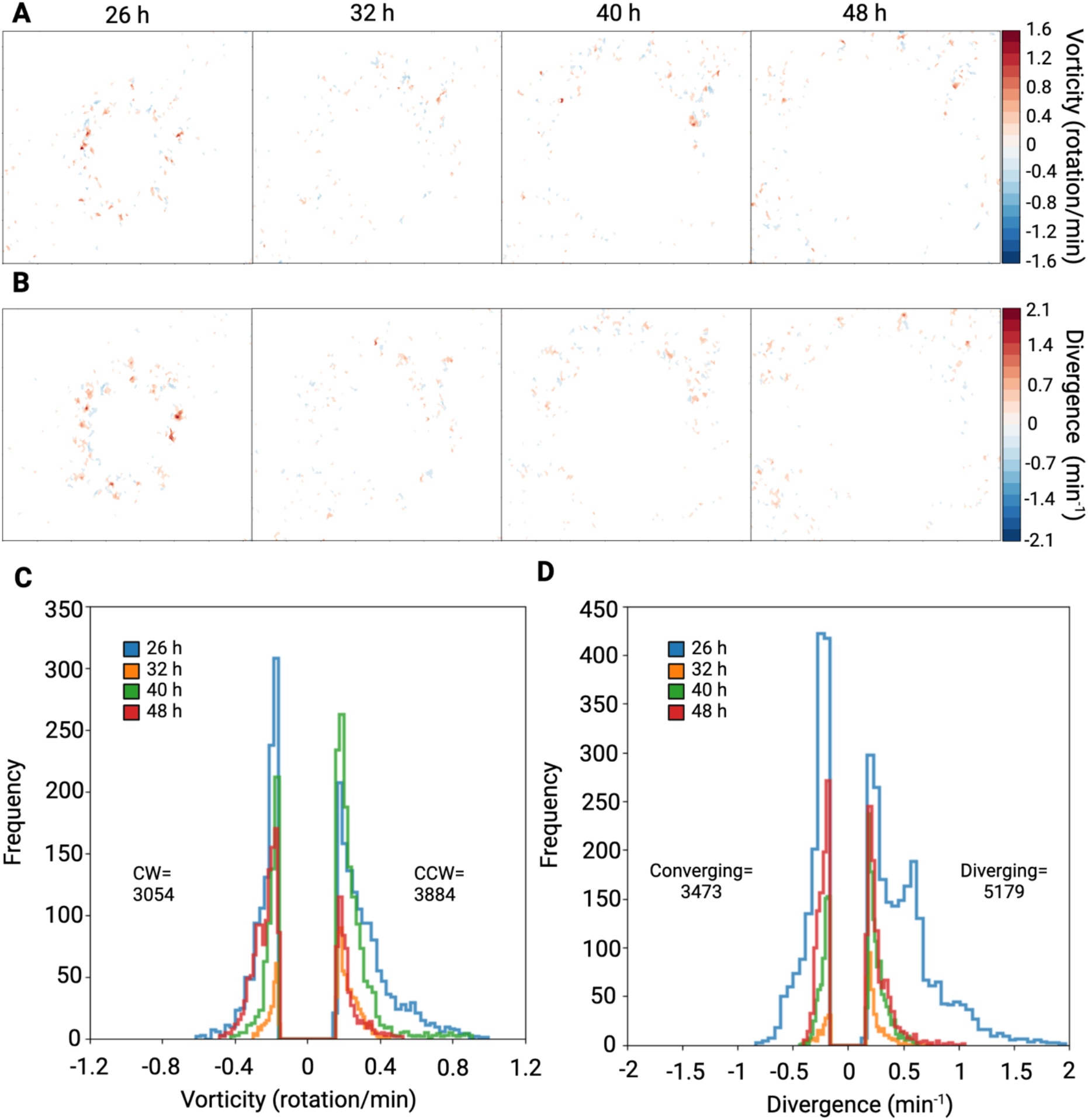
Cell flow derivatives during swarm development. **(A)** Vorticity computed from the velocity maps, with positive vorticity indicating counterclockwise rotation and negative values representing clockwise rotation of swarms, showing highest vorticity initially, and lower rotational movements in later phases. **(B)** Divergence extracted from the velocity map, which quantifies swarmers motion toward or away from a given point in a flow field, with positive values showing divergent movements and negative values indicating convergent movements. **(A, B)** Hotspots of high vorticity and divergence correspond to fireworks and flower-like expansion patterns. **(C)** Frequency distribution of vorticity demonstrating clockwise and counterclockwise rotation, showing slight bias for microswarmers to move in a counterclockwise manner. **(D)** Frequency distribution of divergence, showing greater bias for divergent patterns of microswarmers within the fluid flow.

Around 32 hours, in the areas with the fastest moving swarms, the flower-like patterns divided into two jagged semi-circular shapes. Typically, the semi-circle closer to the periphery expanded outwards, displaying a wave-like pattern of high velocity and low vorticity (**Figure 3B, Figure 4A**). As a wave of swarming bacteria expanded, it merged with another asynchronous wave, forming long and thin wave-like patterns with differing periodicity (**Movie S1**). The CDF shows the range of *v*(*r*) expands from 0.5 µm/min to over 2.5 µm/min, suggesting an increase in *v*(*r*) and possibly in cellular metabolism as well. Notably, a pronounced peak in the CDF between 2 µm/min and 2.5 µm/min at 32 hours suggests that a distinct subset of the cellular population has differentiated itself from the rest of the slower-moving swarm and the faster moving cells exhibit the wave-like motion described above (**Figure 2E**).

By 40 hours, the central region near the initial inoculation spot significantly slowed down, with most motility observed in the periphery. Some of the waves reverted to a more radial-like expansion, with a slight increase in vorticity. Additionally, the swarm velocity at 40 hours resembles the swarm velocity measured at the initial 26 hours **(Figure 2B)**.

Around 48 hours and onwards, the speed of motility substantially decreased, and large dendritic patterns of bacterial growth were observed. This was accompanied by a drop in overall vorticity (**Figure 4A**). Due to the different types of expansion patterns reported here, temporal changes in vorticity were observed, with the maximum vorticity occurring at approximately 1.5 rotations per minute. (**Figure 4A & 4C, S7 & S8**). Our data indicate that swarms vortex in both clockwise and counterclockwise directions. While majority of swarms show bias for counterclockwise vortexing, much like swarms of bacteria using flagella or other means (27, 50, 51). In one case no bias was observed throughout the duration of the experiments, although the signal to noise ratio was higher in this experiment (**Figure 4, S7, S8, Movies S10-S12**). We also found that *C. ochracea* microswarmers commonly exhibit divergent behavior as compared to convergent behavior (**Figures 4D & S7**), but in one case, due to the high signal to noise ratio, similar convergent and diveregent activities were observed (**Figure S8**).

Towards the end of the timelapse, the majority of *v*(*r*) measurements were less than 1 µm/min and scattered microcolonies of *C. ochracea* are observed (**Figure 2B**). In fact, most of the swarms were now non-motile, and due to the swarm patterns described earlier, the overall density of bacteria on the agar plate was unevenly distributed. Interestingly, the regions with higher bacterial concentrations later developed into scattered micro-colonies that were several millimeters to centimeters away and completely disconnected from the parent colony (**Figures 2C-E**). These temporally-driven changes in swarm speed were supported in the additional timelapses generated via this study (**Figure S4 & S5**). Overall, our analysis demonstrate two major modes of movement for *C. ochracea* swarms, one in a wave like pattern where the cells line up and create a linear front, and the second being more radial or fireworks-like expansion where the cells appear to expand outward in all directions while rotating from a central area. This speed and type of movement was not found on 1.5% agar **(Supplemental text, Movies S14-19, Figure S9)**, indicating a specificity of this type of development on softer surfaces.

### Developmental phase transitions are observed within swarms

When the magnitude of *v*(*r*) was plotted as a function of the swarm density corresponding to each *v*(*r*) value, three distinct phases emerged that correspond to swarm propagation and pattern formation at different time points. The significantly motile regions, where the velocity exceeded 0.5 µm/minute, had pixel intensity ranging from 0.3 to 0.7 (**Figure 4A & S6**). At 26 hours, the peak speed was 1.5 µm/minute. At 32 hours, it increased to 2.5 µm/minute and at 40 hours, it dropped back to 1.5 µm/minute. By 48 hours, the speed fell below 1 µm/minute and the pixel intensity ranged from near zero to 0.7.

Additionally, all timepoints displayed an overlap in the 0.5 – 0.7 µm/minute regime, indicative of slow-moving groups that form microcolonies. This behaviour appears to contribute to the floral patterns and isolated swarmers described above for *C. ochracea* swarms. In a contrasting hypothetical scenario, if all cells only slowed around 48 hours to form microcolonies, one might observe a ring-like pattern typical of *E. coli* swarms (48). Hence, the distinct phases, with some overlap, suggest a novel developmental strategy that allows for random placement of the microcolony away from the parent colony, potentially increasing the survival probability of the species.

A numerical agent-based model was developed to simulate cellular dynamics and spatial patterning observed in our experimental data. This model focused on movement, orientation, and collective behavior. The model represents cells as individual entities, each possessing spatial and angular properties in a 2D space. The entities are first positioned within a circle using polar coordinates, with R representing the radius of the circle. As time progresses the positions of the cells transform into Cartesian coordinates within the 2D space. The progression of the simulation is visualized by representing each cell as an oriented rod, which iterates through a series of discrete time steps. The position of the cell changes over time to simulate the motility of the cell.

This model encompasses the position updates for cells in Cartesian coordinates, based on their initial position, velocity, and orientation, with adjustments for local cell density influences on speed and alignment.

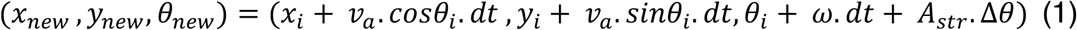

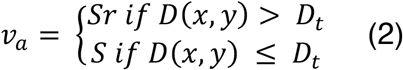

Here, *v* represents the speed of the cell, θ_*i*_ is the initial orientation angle, ω is the rate of change of orientation. *A_str_* is the alignment strength, and Δθ is the angular adjustment based on local alignment. The term *dt* represents the discrete time step. *S* is the standard speed while *Sr* is the reduced speed when the local density exceeds *D_t_*. This numerical model effectively captures the dynamics of cell movement and interaction as they migrate outward from the inoculation spot on a simulated surface. The results from the model demonstrate the emergence of swarm patterns observed in the timelapses (**Figure 5B**). For example, in one field of view from one of the timelapses we can detect nearly all formations that were generated by the simulation (**Figure S5**). Increasing *A_str_* results in less radial or more wavelike patterns, whereas increasing *D_t_* generates swarms with less breaks or protrusions in the swarm pattern, causing more smooth edges. In situations with high *A_str_* and low *D_t_*, non-uniform (i.e. flower petal-like) wavelike pattens are created (**Figure 5B, Movie S13**). High *A_str_* and moderate *D_t_* form incomplete and non-uniform radially expanding swarms. Low *A_str_* and high *D_t_* results in complete radial expansion without many breaks or indentations in the uniform (i.e. continuous curvature) circular swarm. Lastly, low *A_str_* and low *D_t_* results in mostly complete and non-uniform radial expansion with indentations but not many gaps in the swarm front (**Figure 5B**). While our empirical data shows essentially all modes of swarming predicted by the simulation, in the experimental conditions, we typically observe swarms with all *A_str_* values and low to moderate *D_t_* values. We don’t typically observe radially expanding swarms without any breaks or protrusions in the swarm, as predicted in models with high *D_t_*. Hence, the different phases that lead to spatiotemporal patterning of *C. ochracea* swarms depend on changes in cell-density, alignment strength and motile speed.

**Figure 5.**
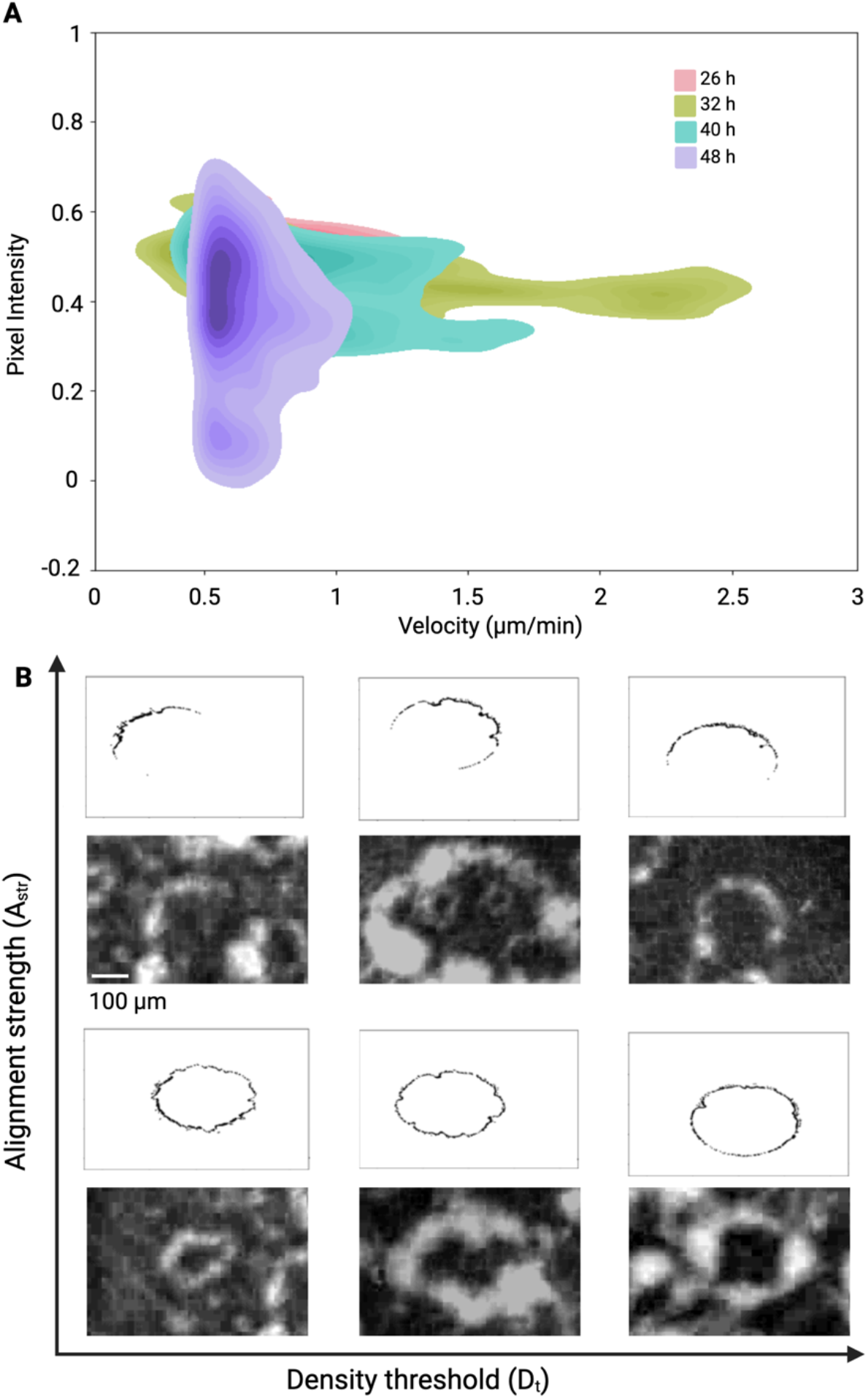
Swarm development model. An agent-based model was developed to characterize microcolony swarm expansion. **(A)** Kernel density (KDE) plots demonstrating different phases during swarm development via changes in velocity and pixel intensity (i.e. cell density). Pixel intensity values between 0.3-0.5 demonstrate the highest speeds at all time points, with the maximum speed achieved at 32 hr. **(B)** Swarm simulations under low and high alignment strengths and density thresholds, and representative images of microcolonies with similar properties to those created in the simulation. Swarms with low alignment strength create connected chains of microcolonies, generating more circular phenotypes than those with high alignment. Swarms with high density threshold create more smooth swarm fronts as compared to lower densities.

## DISCUSSION

Motility is a survival strategy used by organisms across all scales of life. Tracking of multicellular organisms has demonstrated motility as a means of survival and relocation to areas of more optimal living conditions (52). Similarly, fungi release spores, and plants perform seed dispersal to ensure species survival and colonization of new locations (53, 54). This activity is synonymous with bacteria using motility as a means to inhabit new locations. Interestingly, we find that the structure demonstrated by *C. ochraceae* swarms produce dispersed microcolonies which potentially increases the survival probability of the species.

Bacteria have developed diverse motility strategies to relocate and scavenge for nutrients, demonstrating parallels in evolution. The implications of vastly different species evolving machineries that enable their motility demonstrates the importance of active movement in evolutionary fitness. The diverse types of bacterial motility lead to differing expansion strategies and swarm patterns. *Acinetobacter baylyi* and *E. coli* have been shown to create flower-like patterns (24) and *E. coli* are capable of generating chiral flows that are faster than the swarming cells (25). *Myxococcus xanthu*s has been shown to create 3D fruiting body structures under low nutrient conditions (32). *Bacillus subtilis* forms branched, concentric rings (55) that are developed by microscopic whirls and jets (26). *Serratia marcescens* and *Flavobacterium johnsoniae* display vortices that control their resulting swarm phenotypes (27, 56).

Tracking and modeling of swarm characteristics of a common anaerobic oral microbe *C. ochracea* provides novel insight into survival strategies used by bacteria. We find that phase transitions underlie the swarm patterns of coordinated *C. ochracea* cells. These transitions are governed by cell density and alignment strength. The cell density can be increased either through population growth, or through organization of cells via cell surface adhesins. For example, interaction between SprB adhesins on the surfaces of neighboring cells could increase local cell density within micro-swarming populations. Similarly, contact between SprB proteins could create higher alignment strength between cells, causing them to move in a coordinated fashion from physical adherence between cells. External factors could also influence alignment strength of cells, including fluid flows, or neighboring micro-swarmers.

We report that different speed regimes correlate with the types of spatial expansion patterns exhibited by the swarm. Wave-like movements of cells appear to achieve higher speeds than radial, or flower-like swarm patterns. This may indicate two different strategies that underlie swarming. When collectively moving cells exhibit high alignment strength and move along the same axis, they may be able to quickly escape an area. Conversely, utilizing a more radial-type expansion at slower speeds results in cells covering larger surface area, and may demonstrate a scavenging or nutrient seeking phenotype. These observations also align with changes in vorticity over time and we find that swarms of gliding bacteria frequently diverge from the parent swarm. Taken together, these might indicate that swarms of T9SS-driven gliding bacteria display specialized motility tactics in a temporally driven fashion as development proceeds.

We have identified surface stiffness, cell-density, alignment strength, and motile speed as factors that influence spatiotemporal patterning of *C. ochracea* swarms. Future investigation on how metabolism, respiration, chemotaxis, cell division, and polymer production influence spatiotemporal patterning (57–62) might lead to the identification of controllable knobs that can be tweaked to alter the spatial structure of *C. ochracea* swarms. In the human oral microbiota, bacteria from the genus *Capnocytophaga* are found in abundance in supragingival and subgingival biofilms (19). In *in-vitro* settings, non-motile microbes and phages often hitchhike on gliding *Capnocytophaga*, consequently altering the structure of polymicrobial communities. The data presented here show that swarms of *C. ochracea* exhibit spatial patterning, which could potentially serve as a template for seeding the spatial structure of the oral microbiota in a polymicrobial context. Future research exploring the effects of swarming on the spatiotemporal patterning of swarms will not only enhance our understanding of the swarm behavior of gliding *C. ochracea* but will also shed light on its potential role in shaping the structure of oral polymicrobial communities.

## Acknowledgements.

AS was supported by NIH-NIGMS MIRA award R35GM147131. RC and GDD acknowledge support from the Deutsche Forschungsgemeinschaft, grant CO1813/2-1. All figures were created using Biorender.com.

## Author contributions

AZ and AS developed the concept for this study. AZ collected all images and timelapses of developing swarms. RC and GDD performed PIV analysis of developing swarm timelapses on 1% agar. TA and TC captured the three day old images of swarms. WL processed images using support vector machine learning and classification. AS developed the numerical simulations. GDD analyzed developing swarms on 1.5% agar. AZ performed data extraction, analysis and figure preparation for all data collected. AZ, RC, and AS wrote the manuscript .

## Data availability

Custom Python codes for image analysis and example datasets are freely available on our GitHub (https://github.com/amandazdimal/CapnoPIV.git). All other results and data are included in the article, and/or supplemental material.

## Supplementary Text

Macroscopic swarm development was also investigated on TSY with 1.5% agar and 2% agar. However, due to imaging limitations, 2% swarms were omitted from this report due to poor resolution from imaging through the denser agar in these plates. Swarm development on 1.5% agar was tracked using the same method as was described using the 1% agar plates. Similar to the limitation on 2% agar plates, the denser agar and slower movement of swarms made PIV tracking more difficult (**Movies S13-S16**). However, we were able to generate swarm speed from the data collected from PIV. The swarms on 1.5% agar tend to move more slowly, maxing out at speeds around 1.5 μm/min (**Figure S9**), which is 3-times slower than the 4.5 μm/min max speed seen in swarms on 1% agar.

**Supplemental Figure 1.**
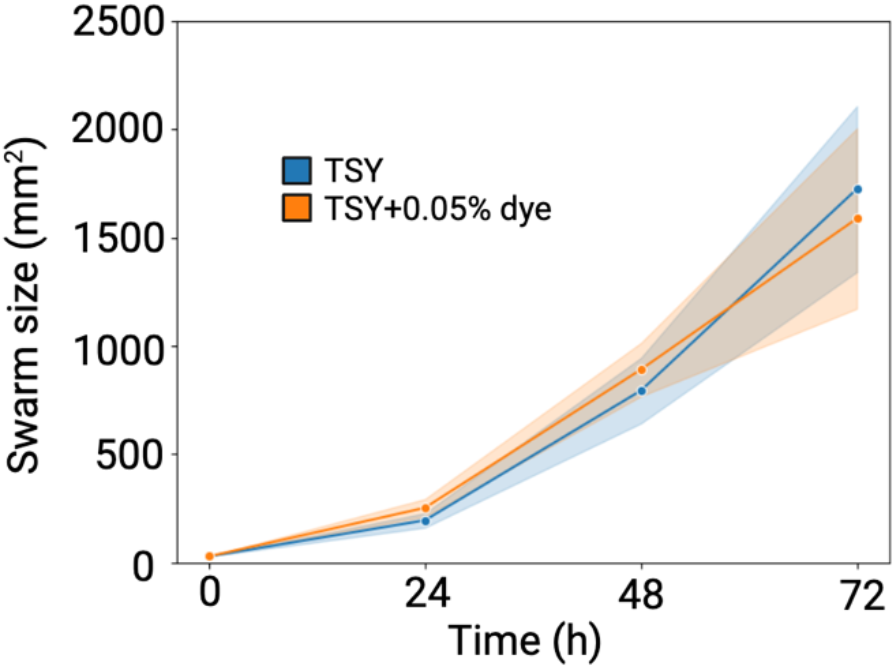
Addition of 0.05% dye on TSY plates with 1% agar show no effect on swarm size. *C. ochracea* swarming was tested on plain TSY with 1% agar, and TSY with 1% agar and 0.05% black food coloring to examine swarming changes in the presence of the dye. There were no statistical differences between conditions for any of the timepoints tested. Data represents 4 biological replicates.

**Supplemental Figure 2.**
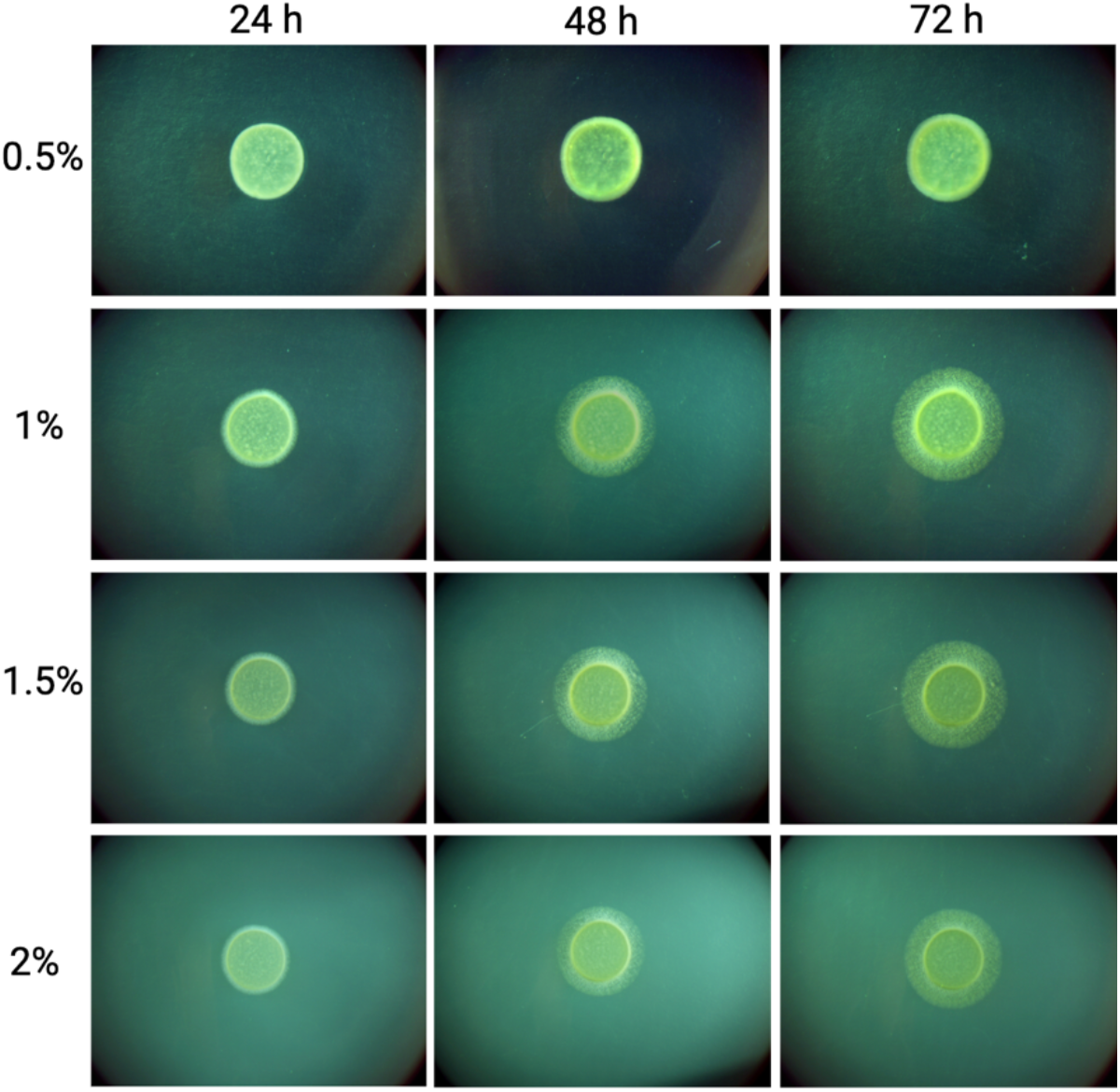
Swarming deficient τιgldK on TSY plates with varying agar concentration. Plates were prepared and inoculated as described in **Figure 1**, but spotted with *C. ochracea* τιgldK cell suspensions. Swarming was not observed, although colony expansion was evident as cells divided and spread out over time.

**Supplementary Figure 3.**
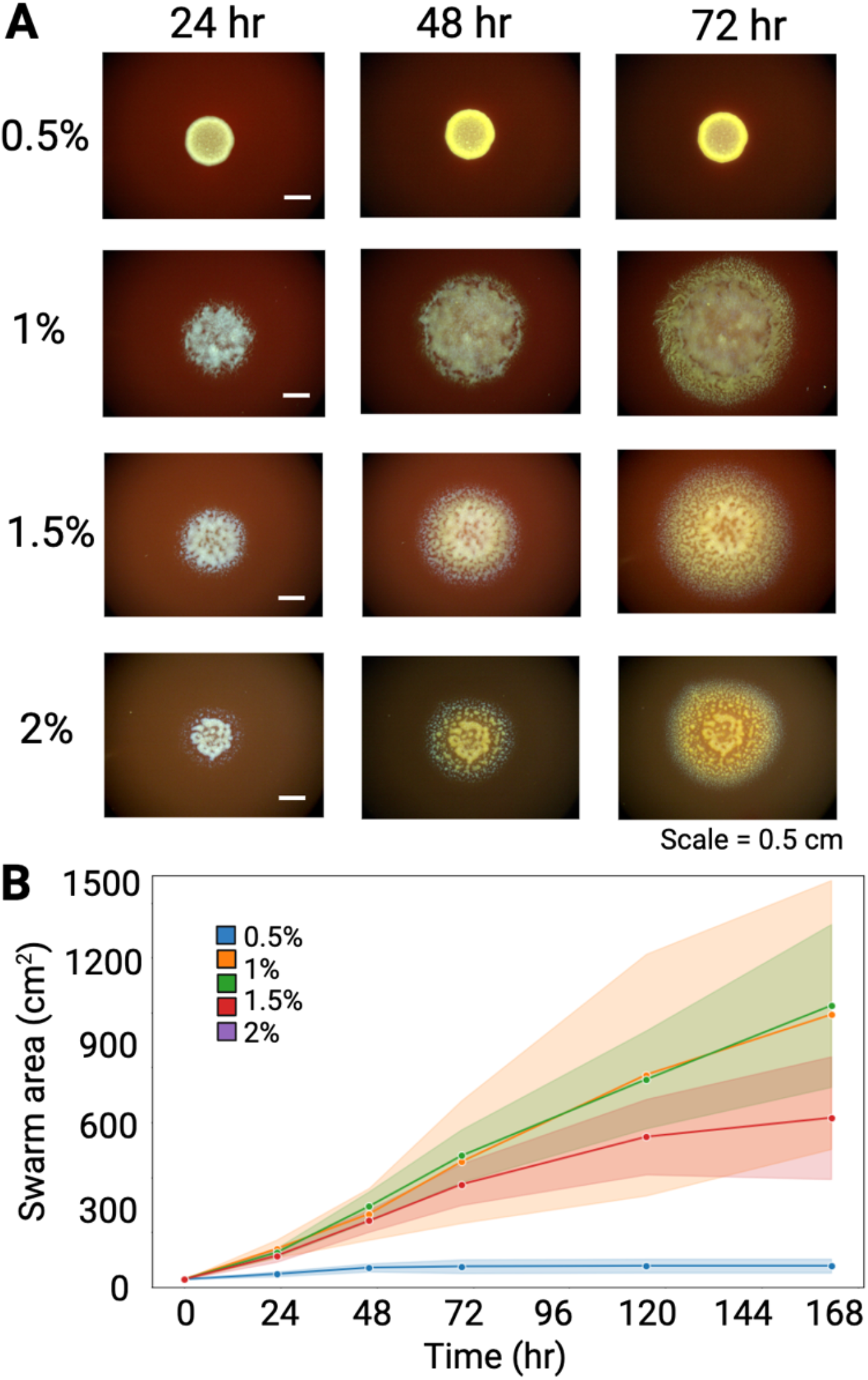
Swarm development on TSY-blood plates. Swarms were tracked as described in **Figure 1**, however these plates contained 5% defibrinated horse blood. **(A)** Photos of *C. ochracea* swarms on TSY-blood plates of varying surface stiffness. **(B)** Swarm area was tracked throughout the duration of the experiment, which showed no statistical significance between 1% and 1.5% agar, unlike results on TSY plates.

**Supplemental Figure 4.**
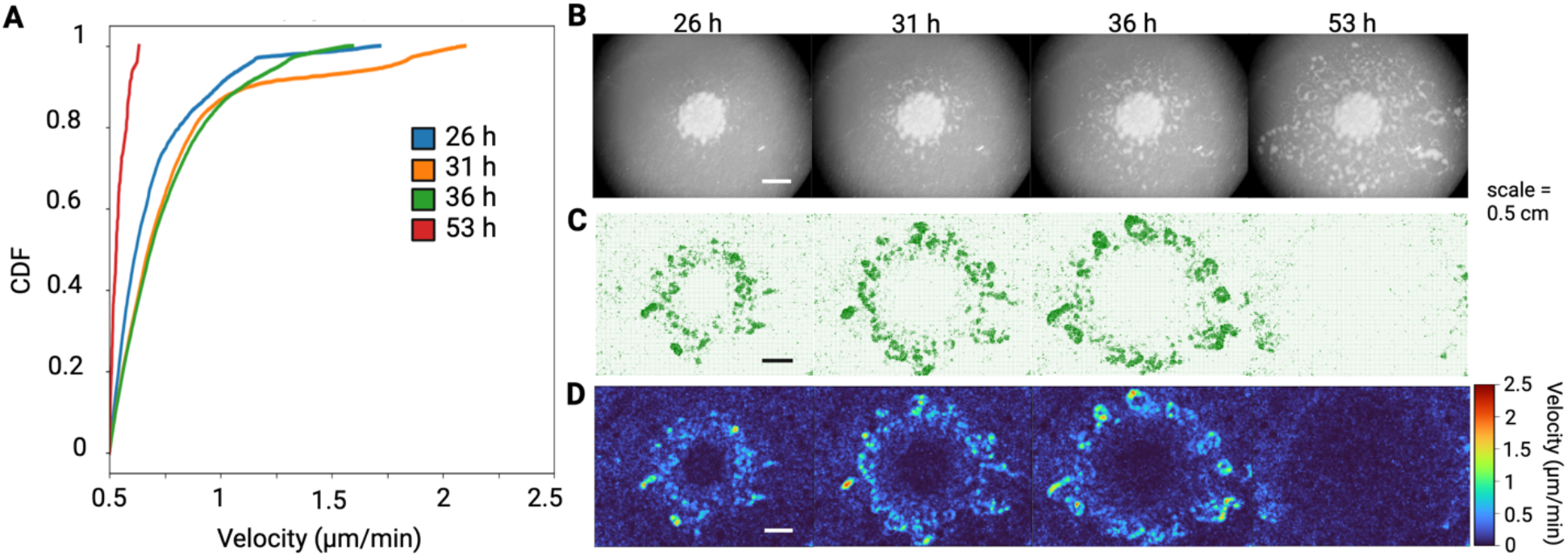
Phase transitions during swarming in timelapse 2. Live imaging of developing swarms were captured as described in **Figure 2**. **(A)** Cumulative density function (CDF) plots for swarm speeds. **(B)** Stereoscope images, **(C)** quiver plots and **(D)** heatmaps of developing swarms during the 4 phases.

**Supplemental Figure 5.**
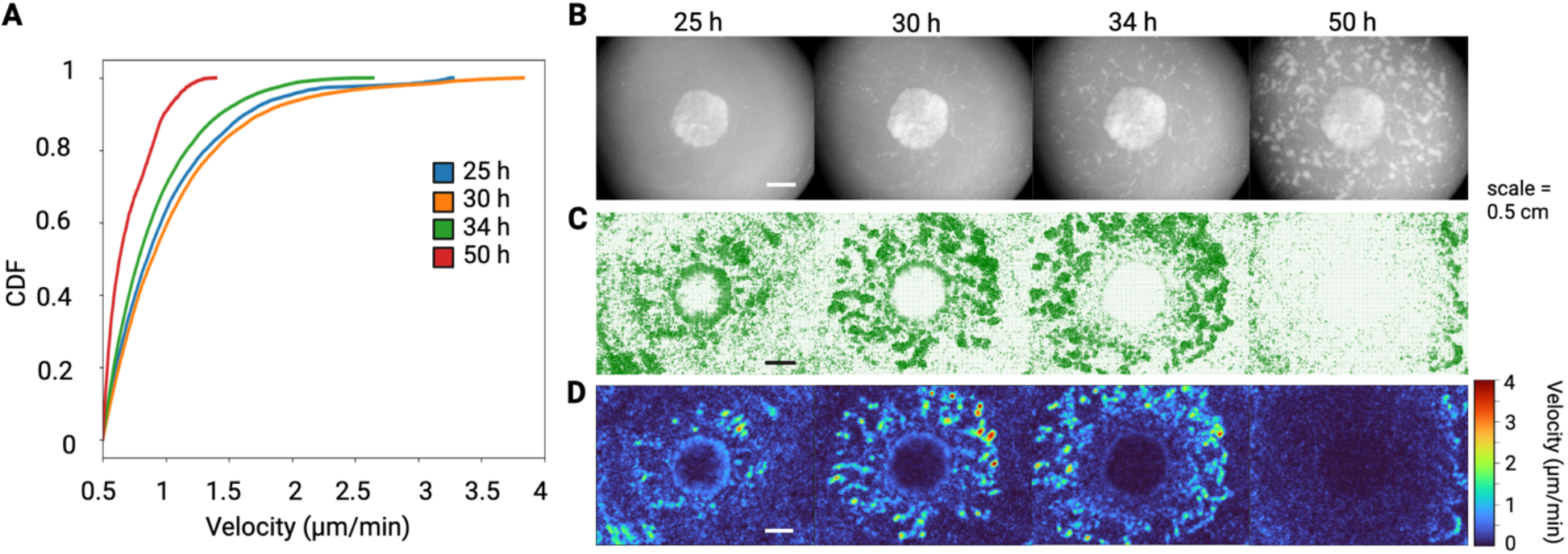
Phase transitions during swarming in timelapse 3. Live imaging of developing swarms were captured as described in **Figure 2**. This timelapse had a different signal to noise profile at the edges as compared with timelapse 1 and timelapse 2. **(A)** Cumulative density function (CDF) plots for swarm speeds. **(B)** Stereoscope images, **(C)** quiver plots and **(D)** heatmaps of developing swarms during the 4 phases.

**Supplemental Figure 6.**
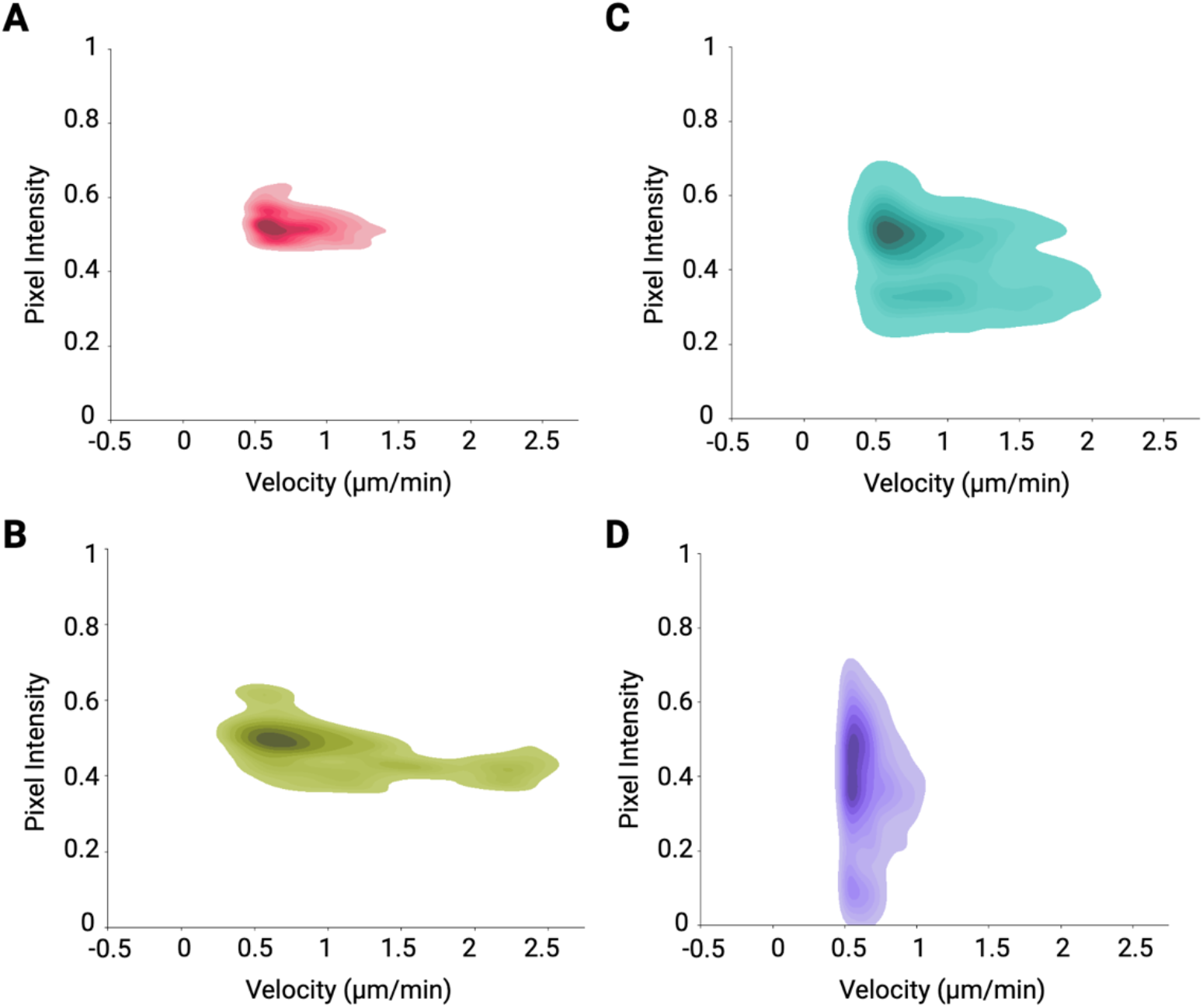
Phase transitions during swarm development. Kernel density (KDE) plots were generated as described in **Figure 4A**. These are the individual KDE plots for the swarm at 26 h **(A)**, 32 h **(B)**, 40 h **(C)**, and 48 hr **(D)**.

**Supplemental Figure 7.**
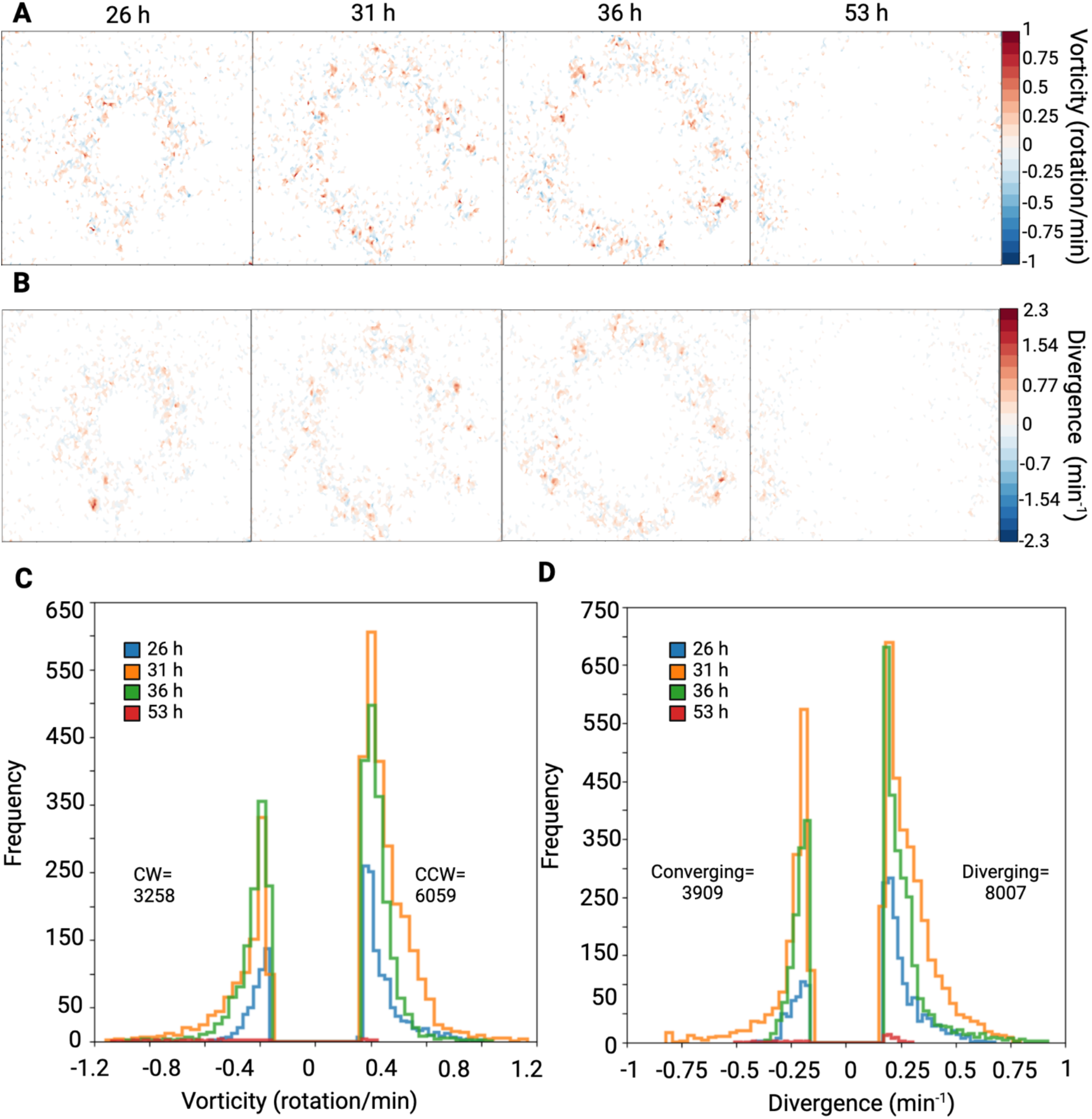
Cell flow derivatives during swarm development, timelapse 2. Velocity maps were used to generate vorticity and divergence data as described in **Figure 4**. **(A)** Vorticity plots showing very similar high vorticity patterns at 31 h and 36 h in phases 2 and 3, with lower rotational movements in early and late phases. **(B)** Divergence plots showing either converging or diverging movements within the flow **(C)** Frequency distribution showing clockwise and counterclockwise rotation, with nearly double the amount of CCW movements vs CW. **(D)** Frequency distribution of divergence, demonstrating significantly more microswarming diverging activity within the flow.

**Supplemental Figure S8.**
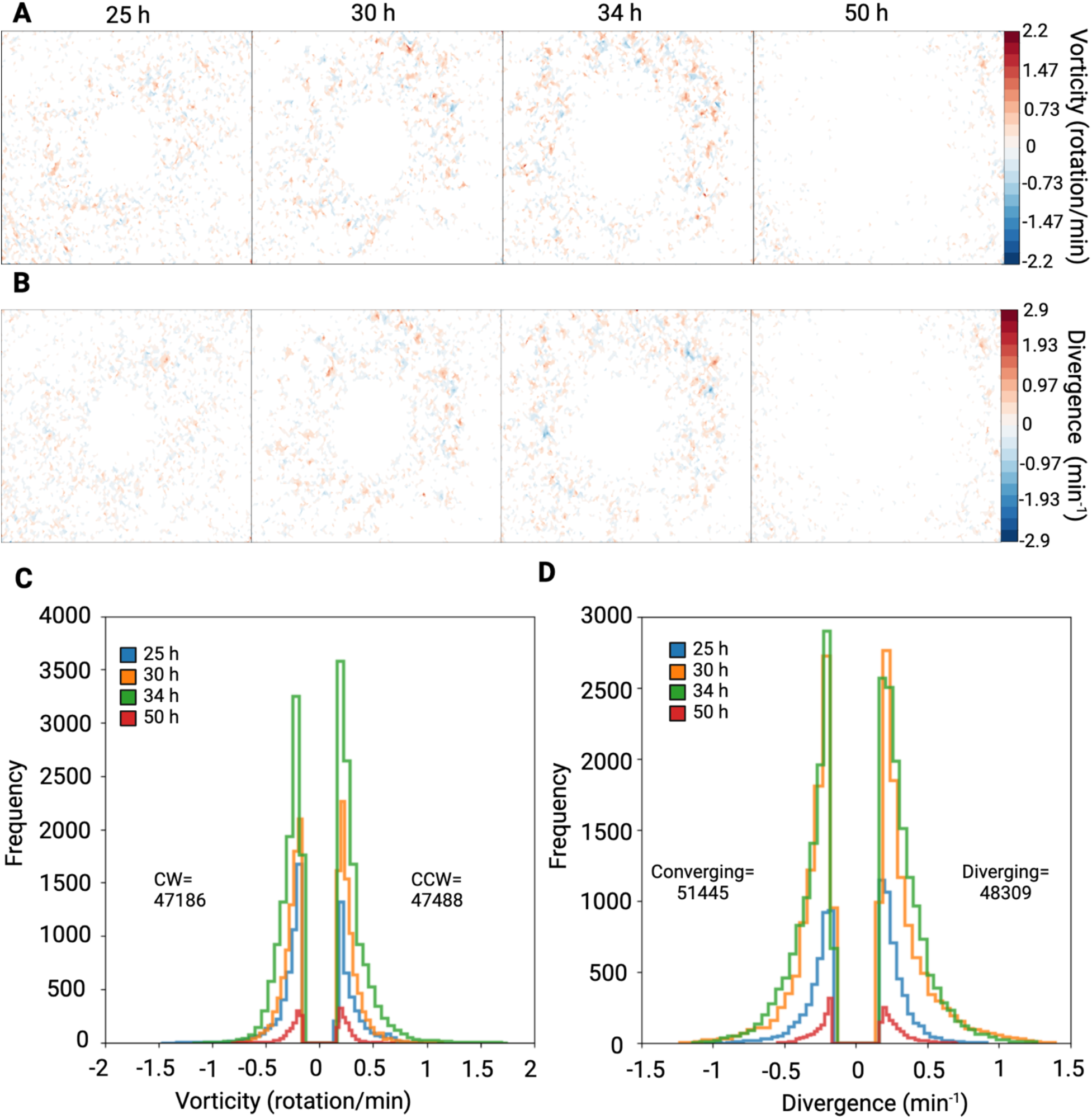
Cell flow derivatives during swarm development, timelapse 3. Velocity maps were used to generate vorticity data as described in **Figure 4**. **(A)** Vorticity plots showing highest vorticity at 30 h and 34 h in phases 2 and 3, and lowest rotational movements in early and late phases, much like **Figure S7**. **(B)** Divergence plots showing a similar amount of converging or diverging movements within the flow, unlike **Figures 4 and S7**. **(C)** Frequency distribution of vorticity demonstrating relatively similar clockwise and counterclockwise rotations. **(D)** Frequency distribution of divergence, demonstrating a slight bias for converging movements, which is in contrast to the majority of diverging activity in timelapses 1 and 2.

**Supplemental Figure S9.**
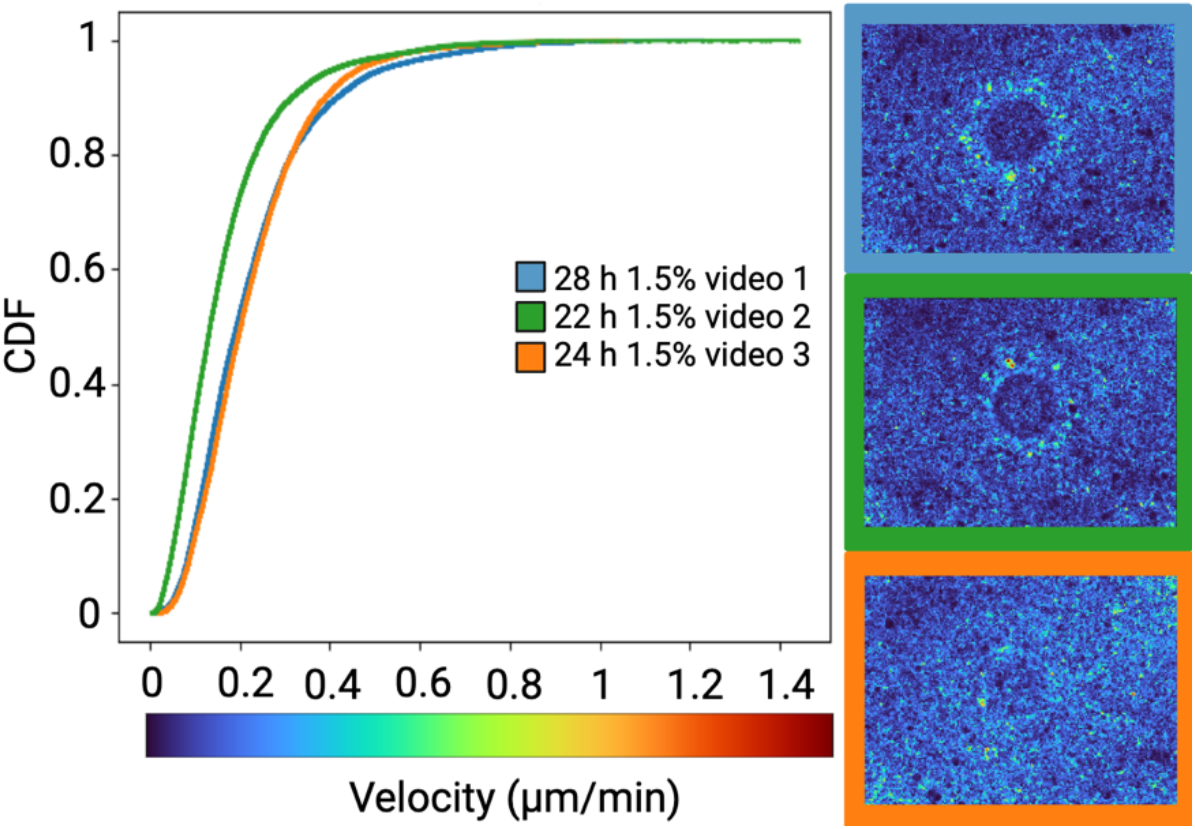
Microswarmer speed on 1.5% agar. Timelapses of developing swarms were captured on 1.5% agar in the same manner for 1% agar. *v*(*r*) was determined for timelapse videos in triplicate. The frames demonstrating the fastest speeds for each timelapse was chosen for CDF analysis. The corresponding heatmaps for the frames are displayed on the right of the plot. Velocities remained below 1.5 um/min in all 3 videos, demonstrating much slower swarm development on 1.5% agar.

**Table S1.**
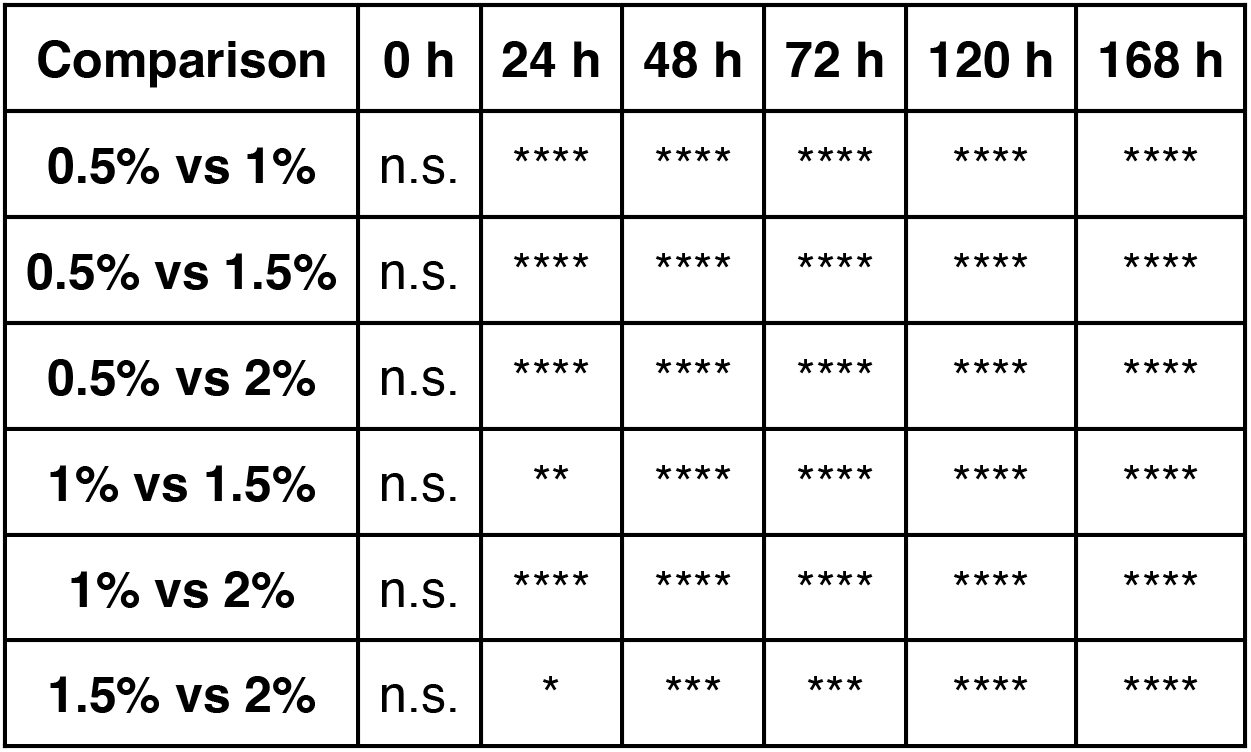
Statistical comparison of *C. ochracea* swarm sizes on TSY agar. . An ANOVA was performed to investigate statistical significance of swarm size at each individual time point. Significance was seen with all comparisons at each time point. (*p ≤ 0.05, ** p ≤ 0.01, *** p ≤ 0.001, **** p ≤ 0.0001).

| Comparison | 0 h | 24 h | 48 h | 72 h | 120 h | 168 h |
| --- | --- | --- | --- | --- | --- | --- |
| **0.5% vs 1%** | n.s. | **** | **** | **** | **** | **** |
| **0.5% vs 1.5%** | n.s. | **** | **** | **** | **** | **** |
| **0.5% vs 2%** | n.s. | **** | **** | **** | **** | **** |
| **1% vs 1.5%** | n.s. | ** | **** | **** | **** | **** |
| **1% vs 2%** | n.s. | **** | **** | **** | **** | **** |
| **1.5% vs 2%** | n.s. | * | *** | *** | **** | **** |

## Supplemetary Movies

**Movie S1.** Timelapse of the developing swarm described in Fig. 2

**Movie S2.** Timelapse 2 of a developing swarm on 1% agar.

**Movie S3.** Timelapse 3 of a developing swarm on 1% agar.

**Movie S4.** PIV timelapse of the swarm in movie S1.

**Movie S5.** PIV timelapse of the swarm in movie S2.

**Movie S6.** PIV timelapse of the swarm in movie S3.

**Movie S7.** Heatmap timelapse of the swarm in movie S1.

**Movie S8.** Heatmap timelapse of the swarm in movie S2.

**Movie S9.** Heatmap timelapse of the swarm in movie S3.

**Movie S10.** Quiver timelapse of the swarm in movie S1.

**Movie S11.** Quiver timelapse of swarm in movie S2.

**Movie S12.** Quiver timelapse of swarm in movie S3.

**Movie S13.** ABM swarm simulation of gliding bacteria.

**Movie S14.** Timelapse 1- PIV of *C. ochracea* swarm development on 1.5% agar.

**Movie S15.** Timelapse 2 PIV tracking *C. ochracea* swarm development on 1.5% agar.

**Movie S16.** Timelapse 3 PIV tracking *C. ochracea* swarm development on 1.5% agar.

**Movie S17.** Timelapse 1 of heatmap plots for developing *C. ochracea* swarms on 1.5% agar.

**Movie S18.** Timelapse 2 of heatmap plots for developing *C. ochracea* swarms on 1.5% agar.

**Movie S19.** Timelapse 3 of heatmap plots for developing *C. ochracea* swarms on 1.5% agar.

## Notes

### Competing Interest Statement

The authors have declared no competing interest.

